# A Deep Operator Network to Predict Thermal Dose on Brain Tumors in MRI-Guided Laser Induced Thermal Therapy: with Synthetic Data

**DOI:** 10.64898/2026.11.17.767113

**Authors:** Yonggang Lu, Max O. Krucoff

## Abstract

**Background:** MRI-guided laser interstitial thermal therapy (LITT) provides minimally invasive option for surgical treatment of recurrent brain tumors. While patient-specific prediction of thermal dose distributions would improve treatment planning, no quantitative methods currently exist for this purpose. Although conventional numerical solvers are being widely investigated, these methods are computationally expensive and often limited by accuracy and reliability.

**Purpose:** Here we propose DeepoLITT, a rapid and accurate model to predict quantitatively heat propagation in LITT procedures. This study represents the first phase of the investigation which focuses on training and validating synthetic datasets to test model’s accuracy, robustness, and computational efficiency.

**Methods:** We built DeepoLITT on a branch–trunk neural network architecture, where the first branch extracts feature from tumor images, the second branch encodes laser parameters (power, duration, and probe orientation), and the trunk models the mapping from spatial coordinates to the resulting temperature field. To train and test the model, we generated synthetic datasets with different noise levels (0∼20%) to mimic LITT procedures with two distinct laser probe types (FullFire and SideFire [Monteris, Inc.]) by a FEM (finite element method) solver. The dataset consisted of 400 samples, of which 90% were used for training and validation and 10% for testing. To evaluate spatial similarity of predicted thermal dose distribution against the ground truth, we employed Dice Similarity Coefficient (DSC) and Hausdorff Distance (HD) computed on CEM43 (Cumulative Equivalent Minutes at 43 °C) isodose contours at thresholds of 2, 10, and 60 minutes. Finally, we evaluated the model’s inference speed with a representative image grid.

**Results:** For the FullFire laser probe, the DeepoLITT model achieved temperature prediction error with average values ranging from 1.16±0.40% to 2.92±0.25%, and peak values ranging from 4.26±1.31% to 14.94±0.43% as data noise increased from 0% to 20%. For the SideFire laser probe, the model achieved temperature prediction error with average values ranging from 3.79±0.53% to 5.13±1.00% and peak values ranging from 15.18±7.15% to 20.75±2.84%. For thermal dose assessment, at the noise-free level for the FullFire laser probe, the model exhibited strong spatial agreement with FEM references, yielding DSCs of 0.96±0.02, 0.97±0.21, and 0.96±0.21 with corresponding HDs of 0.09±0.08 mm, 0.07±0.07 mm, and 0.06±0.08 mm at 2-, 10-, and 60-minute CEM43 thresholds. For the SideFire laser probe, the model achieved DSCs of 0.96±0.02, 0.97±0.21, and 0.96±0.21, with corresponding HDs of 0.20±0.31mm, 0.11±0.30 mm, and 0.15±0.31 mm, respectively. As the noise level increases, these DSC values show a decreasing trend, and the HD values increase. In terms of computational efficiency, the DeepoLITT model inferences temperature maps with 0.04±0.05 seconds per 3D sample ([31 × 31 × 3] grid), compared to 24.32±2.12 seconds required by the FEM solver.

**Conclusions:** The study with synthetic data demonstrates the proposed model is an accurate and rapid method in predicting thermal dose. After testing clinical data, the method has high feasibility to be deployed in real clinical environment.

## 1. Introduction

Laser Induced Thermal Therapy (LITT) is a minimally invasive surgical procedure used to ablate recurrent tumors and radiation necrosis in brain cancer patients ^1–3^. The procedure involves stereotactically placing a laser probe into the tumor in the operating room then transferring the patient with the laser probe into an MRI scanner. The lesion can then be ablated from the MR control room under real-time MR thermometry surveillance, which updates a state space-based temperature measurement every 4-to-8 second to estimate a tissue “kill zone.” In current clinical practice, treatment planning involves choosing a probe trajectory based on empirical prediction of this kill zone, which is basically a ∼1 to 1.5 cm radius from the probe tip (i.e., the maximum estimated range of the treatment based on product limitations). In current systems, the operator can only control the following variables: 1) The anatomical location the laser probe, 2) the type of probe used (i.e., FullFire, which emits a non-directional, roughly spherical heat dose from the tip of the laser, versus SideFire, which emits a directional heat dose out of the side of the laser tip), 3) the number of trajectories used, 4) the on/off time of the laser, 5) the power of the laser (within FDA-approved limits), and 6) the direction of the SideFire probe. There is no established method that can comprehensively incorporate these LITT procedure parameters and thereby accurately predict kill zones, leading to a subjective and highly variable *a priori* predictions of thermal dose and actual ablation volumes.

Classically, the process of heat spread within tumors during laser ablation procedures is physically modeled with optical-thermal partial differential equations (PDEs), which can be solved by numerical techniques (e.g., Finite Element Method [FEM] and Finite Difference Method [FDM]) ^4–8^. However, these approaches rely heavily on precise characterization of laser optics, heat transfer, and tissue properties, each of which can be difficult to obtain in clinical settings. Moreover, these mesh-based methods are computationally intensive and prone to numerical instability, especially under nonlinear conditions commonly encountered during LITT procedures.

Efforts have been made more recently to apply deep learning (DL) and artificial intelligence (AI) methods into scientific and engineering problems. By taking advantage of data driven approaches without discretized meshes and efficient computation of high order derivatives by automatic differentiation, these methods can achieve promising results in resolving spatial-temporal problems ^9–14^. However, conventional DL methods originate from image and language processing, the performance of which is not satisfactory since these methods were not originally designed for resolving physics and engineering problems. Particularly, the high accuracy, precision, speed, and reliability that are required to solve such problems are lacking in conventional DL methods ^13^.

On the other hand, the application of DL specifically designed for scientific applications has exhibited notable effectiveness in addressing complex physics and engineering problems ^15–24^. In mapping input-output parameters in PDEs, neural operator learners show their promise by achieving high prediction accuracy with high precision and computational efficiency. Two canonical approach categories dominate: DeepONet ^19–21^, which represents operators via a branch–trunk network factorization with universal approximation guarantees for nonlinear operators, and Fourier Neural Operator (FNO) ^16–18^, which parameterizes integral kernels in the Fourier domain for efficiency and resolution flexibility. In contrast to classical methods which solve one instance of the equation, both operator learners can learn an entire family of PDEs with better accuracy than previous DL-based solvers under fixed resolution and orders of magnitude faster than numerical PDE solvers ^21^.

Despite the rapid advancement of DL in medical imaging and computational modeling, literature on deep learning - based methods applied specifically to LITT for brain tumors remains limited. Most existing studies have relied on conventional neural architectures such as U-Net ^25^ and long short-term memory (LSTM)^26^ ^27^ networks for tasks like predicting intraoperative temperature fields or estimating regions of thermal damage during LITT procedures. While these models demonstrate the feasibility of learning-based thermal prediction, their accuracy, precision, and computational efficiency have not been thoroughly investigated or quantitatively benchmarked against established ground truth. Moreover, these conventional architectures typically perform direct image-to-image mapping without accounting for procedure-specific variables, including laser power, duration, and probe orientation, that critically influence the spatiotemporal evolution of temperature and thermal dose. Consequently, there remains a significant need for advanced deep learning frameworks capable of capturing the nonlinear and procedure-specific thermal dynamics inherent to LITT while maintaining the computational efficiency required for rapid treatment planning.

In this study, we propose DeepoLITT, a novel deep operator neural network derived from the DeepONet architecture, specifically designed for predicting spatiotemporal heat propagation in pre-operative treatment planning for MRI-guided LITT procedures on brain tumors. This study represents the first phase of the investigation which focuses on training and validating the DeepoLITT model with synthetic datasets to test its accuracy, robustness, and computational efficiency in simulating procedure-specific thermal behavior. The subsequent work will involve integrating clinical data including patient-specific multi-channel MRI images and intraoperative MR thermometry measurements to further enhance the model’s clinical relevance and predictive ability.

## 2. Methods

### 2.1 The Governing Equations in LITT

The LITT thermal ablation process involves complex interactions between laser energy deposition and subsequent heat diffusion within heterogeneous brain tissue. In this study, the procedure is modeled as a coupled radiative transfer and heat conduction problem, formulated through PDEs defined over a spatial-temporal domain. The radiative transfer equation (RTE) governs the propagation and absorption of laser light within tissue, while the Pennes bioheat equation (BHE) describes the resulting thermal diffusion and perfusion-mediated heat loss. This coupled PDE framework captures the nonlinear, dynamic evolution of temperature and thermal dose in response to varying optical and thermal properties ^6,7^.

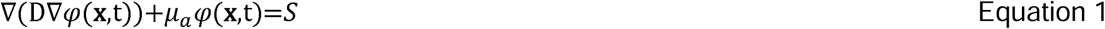

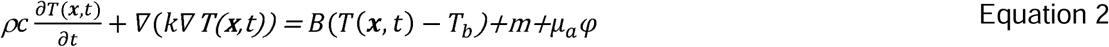

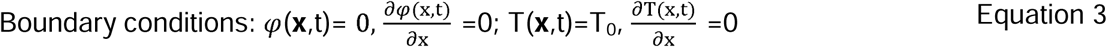

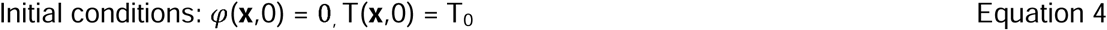

Where φ, optical fluence rate [W/m^2^]; D, optical diffusion coefficient [m]; μ_a_, optical absorption coefficient [1/m]; S, laser light source, P/(AΔz) where P is the laser power [W], A is the area of light emitting element [m^2^], and Δz is the length of light emitting element of laser [m]; T, temperature [K]; T_b_, blood temperature [K]; x, coordinate (x,y,z) [m]; ρ, density [kg/m^3^]; c, specific heat capacity [J/kg/K]; k, thermal conductivity [W/m/K]; =(ωρc)_b_, blood perfusion [W/m^3^K]; m, metabolic heat [W/m^3^].

### 2.2 The Proposed DeepoLITT Method

The above PDEs can be denoted as the following succinct notation ^19,20,28^:

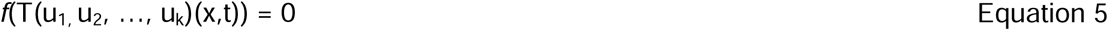

Here f, a symbolic representation of the coupled PDEs; T, temperature field; u_1∼k_, different PDE configurations; (x,t), spatial-temporal domain.

The proposed DeepoLITT method learns an infinite-dimensional function mapping by an operator G_θ_ (θ denotes neural network hyperparameters) that matches the input function space U spanned by the PDE configurations, i=1,2, …, k, to the corresponding function space T spanned by its temperature maps T(u_1,_ u_2_, …, u_k_)(x,t), i.e.,

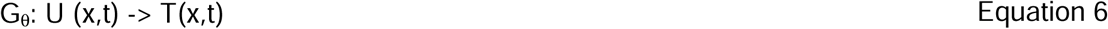

In LITT, u_1∼k_ represents tissue properties, laser source profiles, boundary and initial conditions in spatial-temporal domain. Here we propose a mapping from global scalars (laser power P, duration D, probe orientation O) and image features (I) to temperature maps T.

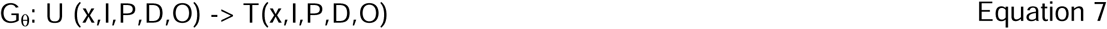

From the universal approximation theorem of operators^19,29^, this mapping can be formulated as a dot product between trainable basis functions (R) and coefficients (B):

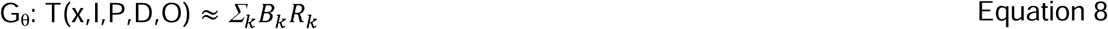

To train the DeepoLITT, a loss function of MSE (Mean Square Error) is optimized to obtain trainable network parameters θ:

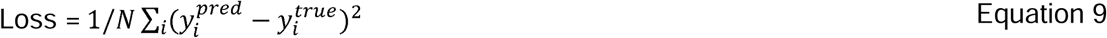

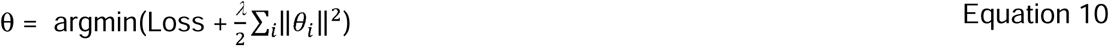

Where N is the number of samples in training, λ is weight decay coefficient.

### 2.3. Network Architecture of the DeepoLITT

We built our model DeepoLITT on a branch-trunk neural network architecture, where a trunk neural network learns the basis function (R) from spatial coordinates, and two branch neural networks learn coefficients (B) from tissue anatomy and laser parameters (Figure 1). Due to established performance of convolution and residue neural networks^30,31^, we built our model on the integration of convolutional and residue blocks. The first branch net comprises three convolutional blocks followed by a series of residual blocks. The convolutional block captures tissue features while repeating residual block is to learn hierarchical representations and complex features of the input data. The second branch and the trunk nets are constructed entirely from stacked residual blocks. Each convolutional block operates over dimensions [input, hidden, latent] and includes convolution, activation, and pooling layers. Each residual block follows the same dimensional structure [input, hidden, latent] but contains additional linear layers to make dimensional projections. Finally, the encoded features from the branch and trunk networks are fused through a fusion net to match the outputs from temperature maps. Through this design, the image branch injects tissue-dependent fields into the network, allowing the same trunk basis to adapt to different shapes and properties while the laser branch modulates the magnitude and temporal duty of heating.

**Figure 1.**
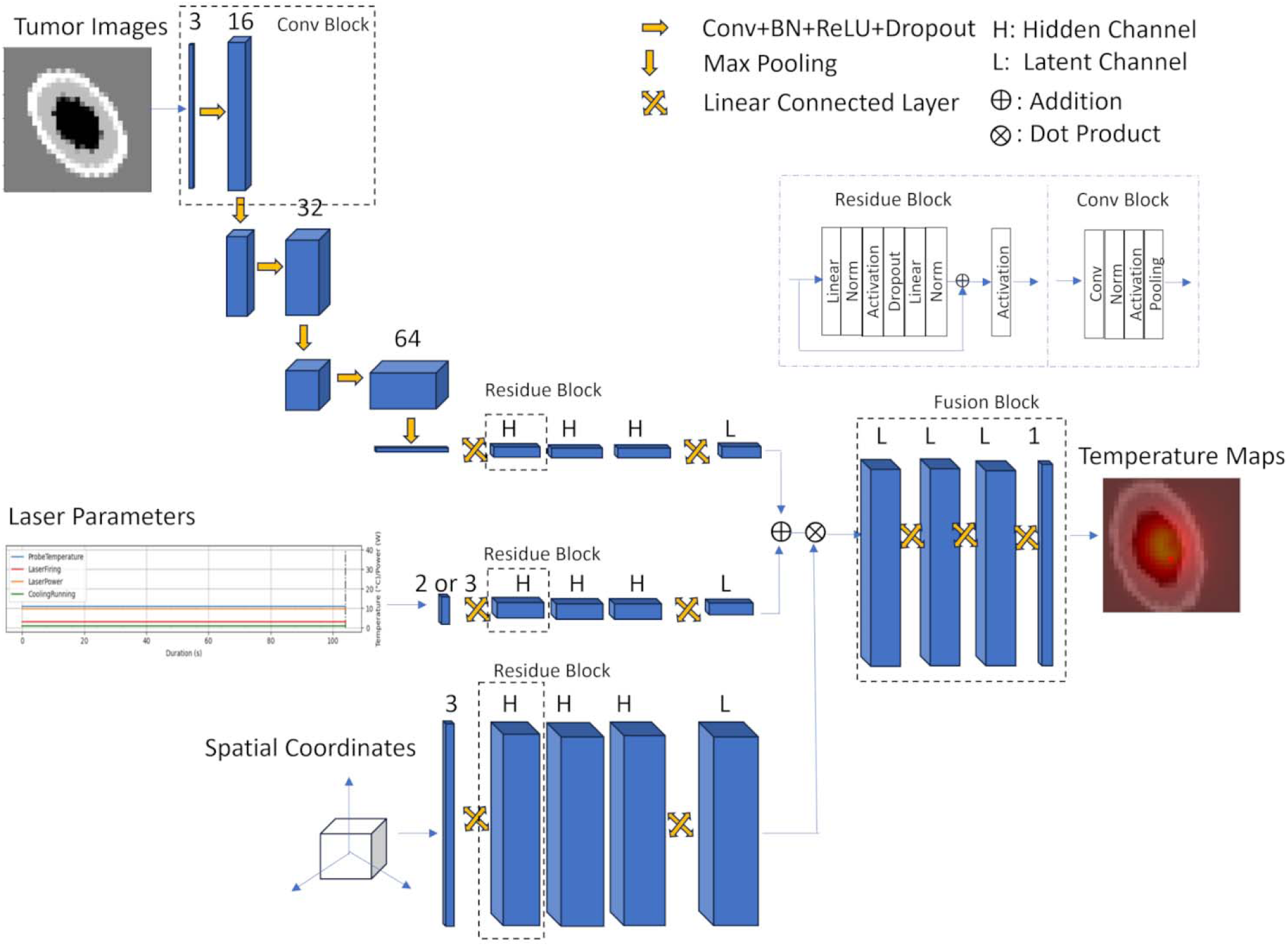
The DeepoLITT network architecture. The number above each layer block indicates the number of channels in that layer.

To identify an optimal model architecture, we performed an automated search procedure using the Optuna framework^32^. The optimization explores diverse combinations of architectural components and hyperparameters through randomized trials. Leveraging the Tree-structured Parzen Estimator (TPE) algorithm^33^, a Bayesian sampler models the predefined hyperparameter space and adaptively prioritizes regions with higher potential performance.

The predefined search space for architectural components were as follows: (i) The structure of branch #2 with choices between a single branch for grouped laser parameters (PDO) versus separate branches for each individual parameter (P,D, and O); (ii) The fusion mechanism, comparing a simple concatenation layer with a dedicated fusion block; and (iii) The normalization strategy, selecting between layer and batch normalization. The hyperparameter search space was defined from the following: latent dimension [8, 128], hidden dimension [8, 256], optimizer type [Adam, AdamW, RMSprop], learning rate [0, 8E-4], number of convolutional blocks [0, 5], and weight decay [0, 8E-5].

### 2.4 Synthetic Datasets Generated by the FEM solver

The coupled RTE-BHE equation forms the physical foundation for generating reference data to train and test the DeepoLITT model. In this study, we used a FEM solver to resolve the coupled equations to create synthetic data as ground truth for model training and testing.

To simulate tumor tissue appearance in MRI T1-weighted contrast-enhanced images, a 3D volumetric digital phantom with dimensions of 60 × 60 × 60 mm³ was created as a computational domain illustrated in Figure 2. The volume comprises a tumor region, including a tumor core, necrotic core, and tumor rim, embedded within normal brain tissue, labeled as 1, 2, 3, and 0, respectively. Each tissue type is assigned distinct image intensity and biophysical properties. To generate a diverse synthetic dataset, this base volume image is augmented by introducing random variations in intensity, size, rotation, translation, scaling, and shear. Specifically, intensity values are randomly sampled within the following ranges: tumor core [0.8,0.9], necrotic core [0.9,1.0], tumor rim [0.5,0.6], and normal tissue [0.6,0.7]. The tumor core radius is varied between [15, 20] mm, the necrotic core radius between [8, 10] mm, and the tumor rim thickness between [3,8] mm. Geometric transformations include rotation angles from –20° to 20°, scaling factors from 0.5 to 1.5, translations from –5 to 5 mm, and shear values from –0.5 to 0.5. The optical and thermal tissue properties, including absorption coefficient, scattering coefficient, thermal conductivity, specific heat, blood perfusion, and metabolic heat generation, for each tissue type are adopted from Table 1 ^34–39^.

**Figure 2.**
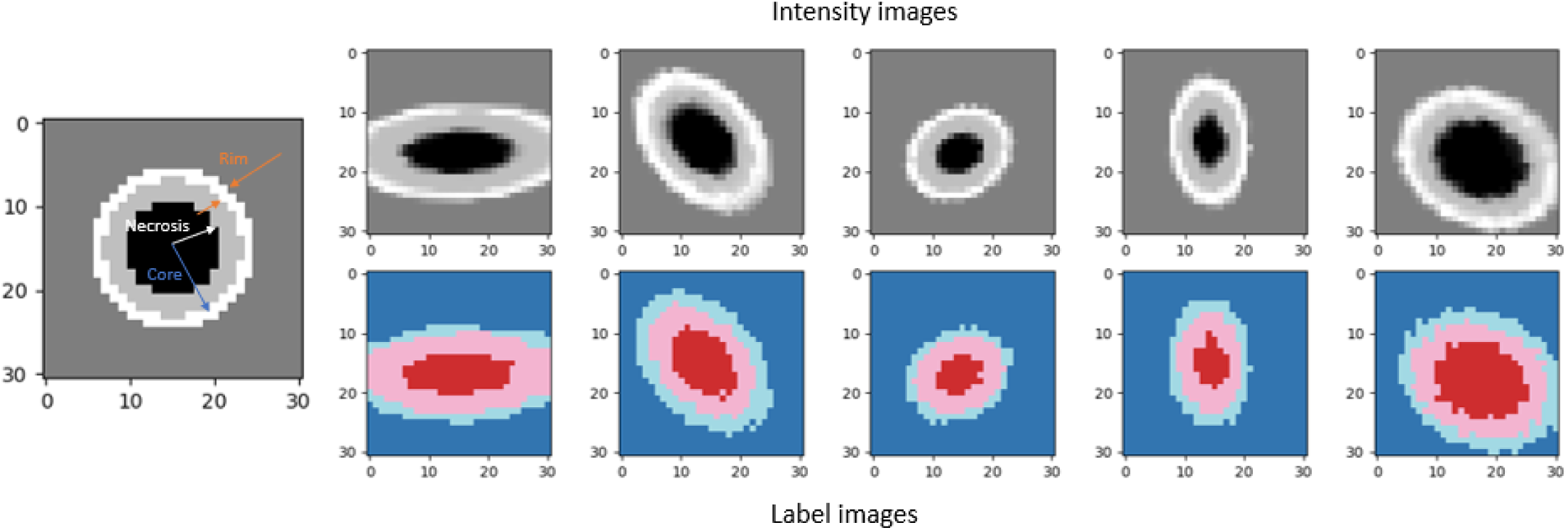
Synthetic tumor tissue images with labels. The top row shows intensity images, and the bottom row displays corresponding label images. From left to right, the panels illustrate the base image followed by five selected randomly augmented samples.

**Table 1.** Tissue optical and thermal properties.

| Tissues/ properties |  | Optical absorption coefficient (1/m) | Optical scattering coefficient (1/m) | Thermal conductivity (W/m.K) | Density (kg/m<sup>3</sup>) | Specific heat (J/kg.K) | Blood perfusion (W/m<sup>3</sup>) | Metabolic heat (W/m<sup>3</sup>) |
|---|---|---|---|---|---|---|---|---|
| Normal brain tissue |  | 50 | 5700 | 0.53 | 1045 | 3630 | 32482 | 17000 |
| Tumor | Core | 60 | 5500 | 0.45 | 1025 | 3300 | 37440 | 20000 |
|  | Rim | 70 | 6000 | 0.48 | 1035 | 3700 | 40000 | 30000 |
|  | Necrotic | 90 | 9000 | 0.35 | 1010 | 3500 | 0 | 0 |

The simulation parameters are based on the specifications of the NeuroBlate System (Monteris Medical, Winnipeg, Canada) ^40^, which is clinically employed at our institution for MRI-guided LITT treatment for brain tumors. A laser probe with a diameter of 3.3 mm and an embedded light-emitting element is modeled at the center of the domain. For the FullFire probe, the emitter is represented as a line source with a ra ius of 1 mm and length of 6 mm, whereas for the SideFire probe, the light emission is modeled as a focal region with a 1 mm radius, transmitting through a rectangular slit window on the probe surface with dimensions of 0.4 × 0.7 mm² at an angle of 78° relative to the probe axis. Laser heating is simulated with power levels randomly sampled from [8, 15] W, exposure durations from [100, 300] s, using a temporal resolution of 8 s, and probe angular orientations from [-180°,180°] (for the SideFire probe). Figure 3 illustrates the detailed geometries of probe structure and light emission angle used in the simulations.

**Figure 3.**
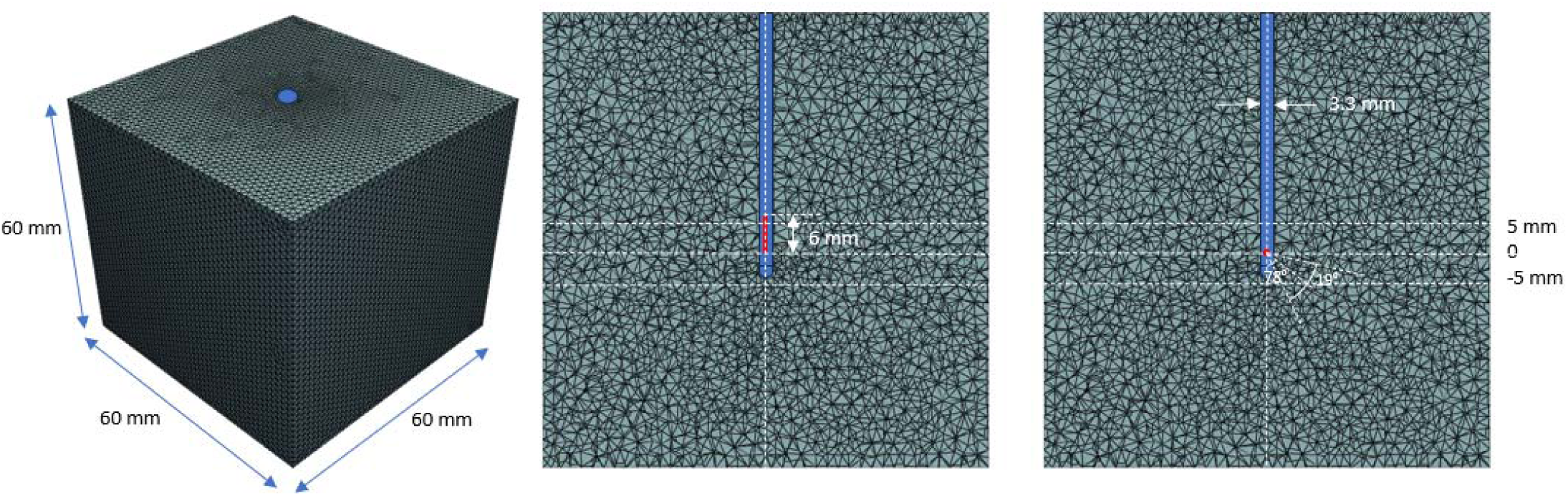
The computational domain with FEM (Left: a cubic volume containing tumor tissues; middle: a FullFire laser probe in the domain; right: a SideFire laser probe in the domain). The coordinates along the probe trajectory at [-5, 0, 5] mm denote the three image locations where the computed temperature fields are sampled.

The coupled radiative and heat transfer PDEs were numerically solved within a FEM framework implemented in FEniCS/DOLFINx^41^, where continuous function spaces were defined for the fluence and temperature fields, while cell-wise function spaces were used for spatially varying tissue properties. The initial conditions (time = 0 s) were set to zero light fluence and baseline temperature of 35 °C, representing physiological equilibrium. Boundary conditions at the domain edges were maintained at the same temperature while boundary temperature at the probe surface was set to be probe cooling temperature which is 11 °C for the FullFire probe and 8 °C for the SideFire probe. The computational domain was discretized using a tetrahedral mesh with element sizes ranging from 0.5–1.5 mm, resulting in 56,818 nodes and 342,869 elements. A direct LU decomposition solver was employed to obtain numerical solutions for the light fluence and temperature fields.

To emulate MR thermometry outputs from the NeuroBlate system, three 2D image slices were sampled at axial positions of –5 mm, 0 mm, and +5 mm along the probe shaft, each with an in-plane resolution of 128 × 128 pixels and spatial resolution of 2 × 2 mm². The simulated temperature maps were normalized to a range of 35–100 °C to avoid exploding gradients and subsequently imported into the DeepoLITT framework as ground-truth for model training, validation and testing.

Four hundred randomizations were applied to the datasets for both the FullFire and SideFire probes, resulting in augmented datasets with 15,090 time points for the FullFire probe and 15,370 time point for the SideFire probe. Ten percent of each dataset was held out for testing, and the remaining data (90%) was used for training and validation.

### 2.5 Model Training, Validation and Testing

A k-fold cross-validation ensemble method (k = 5) was used to train and validate the model. The training dataset was randomly partitioned into five equal subsets; in each round, one-fold was held out for validation while the model is trained on the remaining four folds. This process yielded an ensemble of five trained models, and performance was reported as mean ± standard deviation across folds. As mentioned before, the unseen 10% samples of the dataset were held out for independent testing.

The optimizer with initial learning rate and weight decay was selected from the optimization searching with incorporated strategies of early stopping, learning-rate scheduler, and loss function regularization. The learning rate scheduler automatically adjusts the initial learning rate when the loss function reaches a plateau or starts to increase. To refine the model parameters, the above outputs were further optimized using a deterministic L-BFGS optimizer with a full batch data.

To assess model robustness, the dataset was augmented with varying levels of Gaussian noise, where the noise level is defined as the ratio of standard deviation to signal mean (σ/μ) taking values of [0, 5%, 10%, 15%, 20%]. Noise was applied to both the tissue images and temperature maps.

To evaluate temperature prediction errors of the model with the testing dataset, the following error metrics were defined:

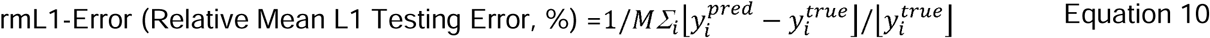

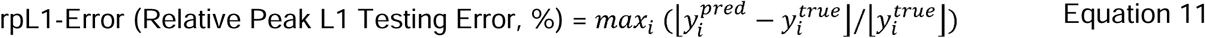

Where M is the number of testing samples.

To estimate correlation between predicted and true values, a linear regression was analyzed, yielding slope, intercept, and R² score.

To illustrate the model performance visually, a parity plot which displays correspondence between predicted versus true values across all evaluated voxels (a fold in this study), an error histogram which quantifies the voxel-wise distribution of prediction errors, and a sample-wise comparison plot which compares the predicted and true temperature values across samples (to improve visual clarity, only 1,000 samples were plotted) were displayed. Especially, the sample-wise comparison plot was illustrated with a 95% confidence interval (mean ± 1.96 standard deviation). Additionally, temperature maps (a 3D volume image and a 2D image in the middle slice) of ground truth, prediction, and their absolute errors from a random sample were further visualized and compared.

The standard thermal dose metric CEM43 (Cumulative Equivalent Minutes at 43°C) iso-dose contours at 2, 10, and 60 minutes ^42–44^ were used to evaluate thermal dose distribution on the target. CEM43 iso-dose contours at 2-10 minutes represent early heat accumulation while CEM43 iso-dose contours at 10 - 60 minutes reveal prolonged thermal exposure. Similarity of CEM43 = 2, 10, 60 iso-dose contours between the DeepoLITT model and the ground truth were evaluated using average Dice Similarity Coefficient (DSC) and Hausdorff Distance (HD) along temporal course.

All experiments were implemented using PyTorch v2.7.1 with CUDA 12.8 support for GPU acceleration and executed on a Linux workstation equipped with an AMD processor and an NVIDIA A10 GPU.

## 3. Results

### 3.1 Architecture Ablation and Optimization

The architecture ablation study (Figure 4) demonstrated that the network with a configuration of a grouped laser parameter branch (the PD branch choice for FullFire, the PDO branch choice for SideFire), a layer normalization, and a linear fusion block had the best performance among the networks with other configurations. Specifically, a branch which incorporates grouped power–duration (PD) for the FullFire probe, or power–duration-orientation parameter (PDO) for the SideFire probe, outperformed separate laser parameter branches (P_D for FullFire or P_D_O for SideFire); the network which add a linear fusion block instead of a simple concatenate layer yielded lower validation loss; the network which uses a layer normalization rather than batch normalization reduced validation loss in model performance.

**Figure 4.**
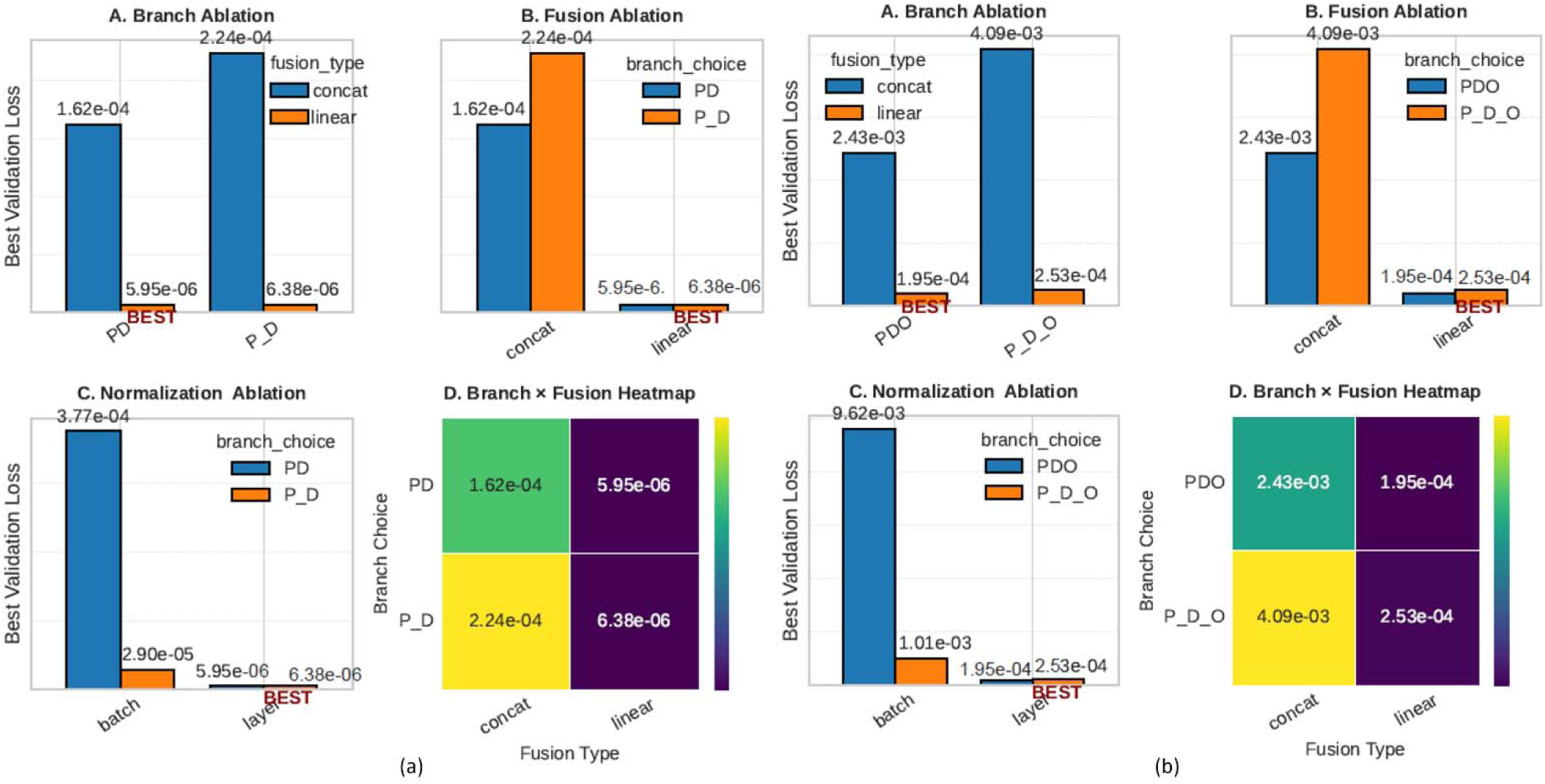
Architecture ablation. (a) for the FullFire probe (PD or P_D denotes grouped or separate power and duration branches); (b) for the SideFire probe (PDO or P_D_O denotes grouped or separate power, duration and orientation branches). BESTs in the bar plots denote the best network configurations searched from the Optuna algorithm. The color bars in heatmaps denote the value range of validation loss.

From the analysis of parameter importance (Figure 5), the FullFire and SideFire probes showed different hyperparameter preferences. For the FullFire probe, the learning rate, latent dimension, dropout rate, hidden dimension, branch choice, and fusion type emerged as the primary influential hyperparameters affecting model performance. The optimal set of hyperparameters identified through the optimization process for the FulFire probe were as follows: latent dimension = 22, hidden dimension = 254, optimizer = AdamW, convolutional blocks = 2, dropout rate = 0.07, learning rate = 3.1 E-4, and weight decay = 9.1 E-5. On the other hand, for the SideFire probe, the fusion type, number of blocks, hidden dimension, learning rate and dropout rate are shown as the primary influential hyperparameters affecting model performance. The optimal set of hyperparameters identified through the optimization process for the SideFire probe were as follows: latent dimension = 46, hidden dimension = 279, optimizer = AdamW, convolutional blocks = 3, dropout rate = 0.07, learning rate (lr) = 5.5 E-4, and weight decay = 9.11 E-5. These optimized configurations were subsequently used for model training and testing in experiments. With these optimized network architectures and hyperparameters, in training a k-fold (k=5) cross-validation ensemble model, approximate 3 hours were spent for the FullFire probe model and 4 hours are spent for the SideFire probe model.

**Figure 5.**
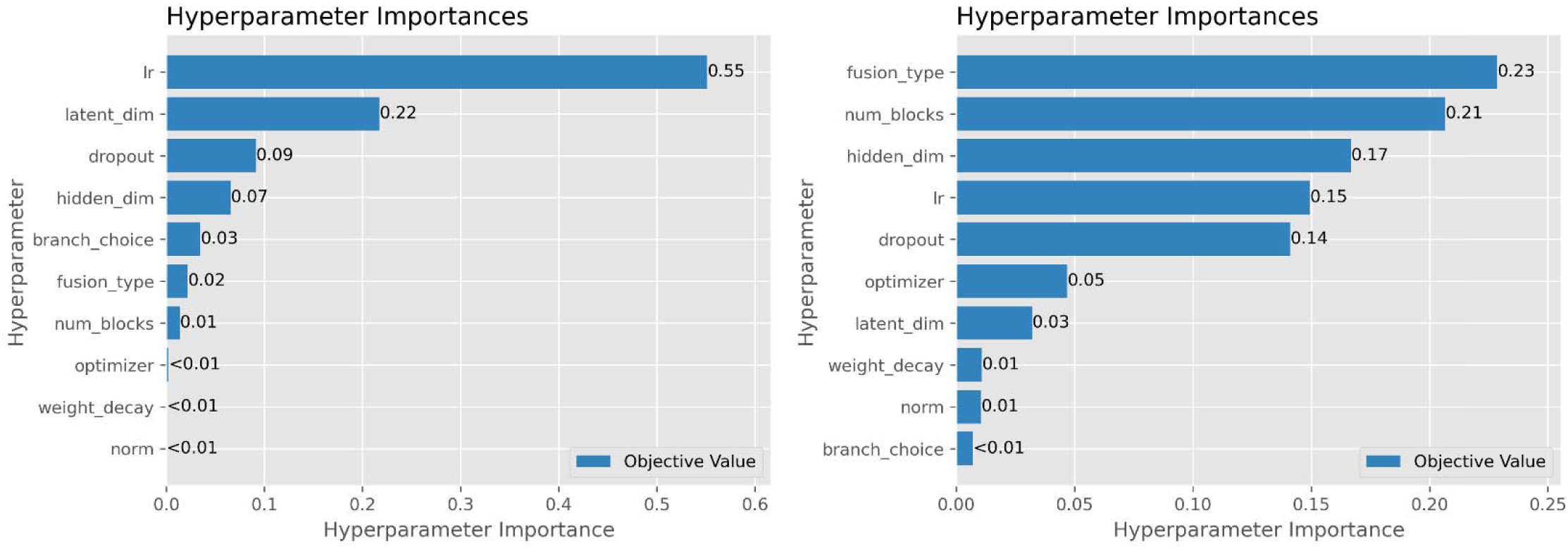
Hyperparameter importances. Each bar magnitude denotes its objective value in optimization, representing relative importance of each hyperparameter. Left: for the FullFire probe; Right: for the SideFire probe.

### 3.2 Prediction Error of Temperature

To evaluate generalizability of our model, we tested the performance of temperature inferencing with the testing data. The performance of the DeepoLITT model, evaluated on testing dataset across different noise levels, is illustrated in Figures 6 (the FullFire laser beam) and 7 (the SideFire laser beam), with examples shown for noise levels at 0 and 20%. The model exhibited strong linearity between predicted and true temperature values, with regression slopes approaching unity, indicating accurate prediction and calibration. Error histograms reveal only a small number of voxels with large deviations, while the confidence intervals remain narrow even under high noise conditions, reflecting the model’s robustness to data perturbations. The predicted temperature maps from randomly selected test samples closely match the ground truth at low noise levels, with discrepancies becoming more noticeable as noise increases. Importantly, the spatial shapes and structural patterns of the temperature distributions are well preserved, suggesting that the model effectively learns the underlying physics governing heat diffusion and successfully captures the tissue-specific thermal behavior.

**Figure 6.**
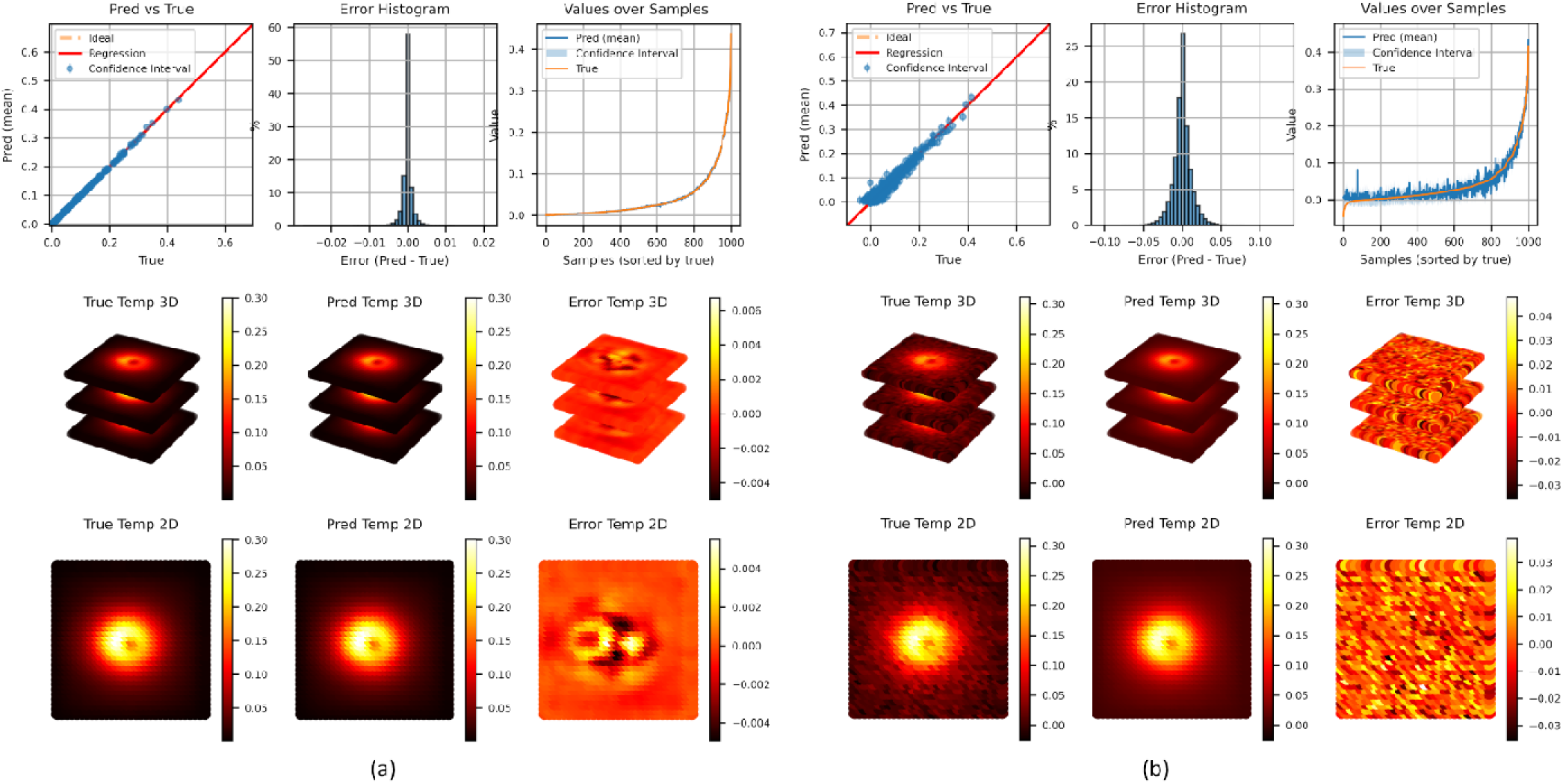
Temperature prediction error of the DeepoLITT model for the FullFire laser beam at noise level =0 (a) and = 20% (b) with the testing dataset. For each noise level, the top panel displays plots of parity, error histogram and confidence interval of samples, the middle and bottom panels showcase a 3D volume mage and its middle plane image, respectively.

**Figure 7.**
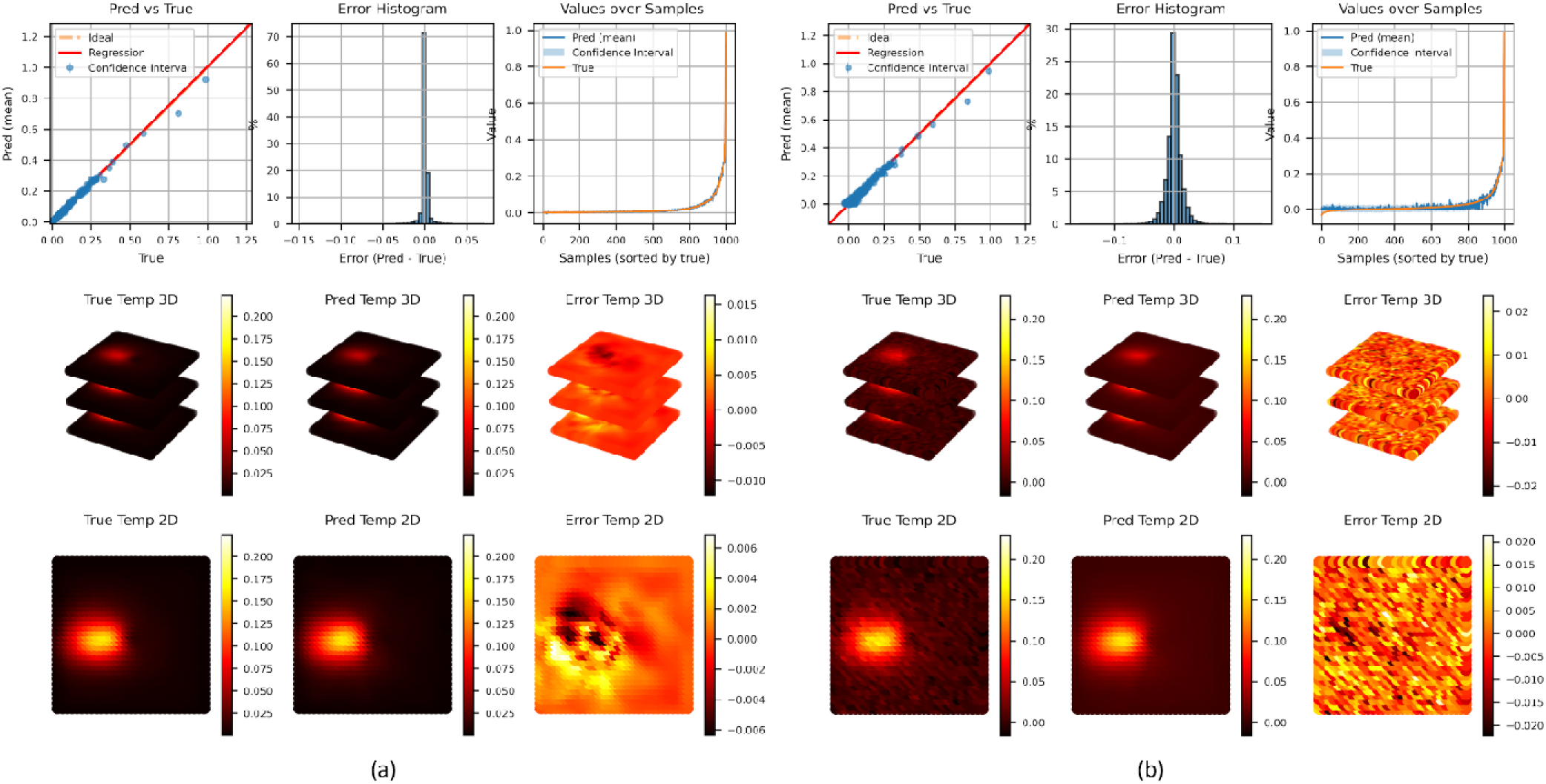
Temperature prediction error of the DeepoLITT model for the SideFire laser beam at noise level = 0 (a) and = 20% (b) with the testing dataset. For each noise level, the top panel displays plots of parity, error histogram and conformal band of samples, the middle and bottom panels showcase a 3D volume image and its middle plane image, respectively.

Tables 2 summarizes the effect of different input noise levels on the predictive performance of the model. As the noise level increase from 0 to 20%, a gradual degradation in accuracy is observed a ross all error metrics (rmL1-Error %, and rpL1-Error %). For example, under noise-free conditions, the model achieves near-perfect agreement with the FEM reference, with an average L1-Error of 1.16±0.40 % (FullFire) and 3.79±0.53 % (SideFire). However, when the noise level is further raised to 20%, performance degradation becomes more pronounced with an average L1-Error of 2.92±0.25 % (FullFire) and 5.13±1.00 % (SideFire).

**Table 2.** Temperature prediction errors.

| Noise level | rmL1-Error (%) |  | rpL1-Error (%) |  |
|---|---|---|---|---|
|  | FullFire | SideFire | FullFire | SideFire |
| 0 | 1.16±0.40 | 3.79±0.53 | 4.26±1.31 | 15.18±7.15 |
| 5% | 1.23±0.17 | 3.99±1.23 | 5.37±0.56 | 15.40±3.67 |
| 10% | 1.89±0.23 | 4.07±2.26 | 8.82±0.51 | 15.78±1.64 |
| 15% | 2.39±0.18 | 5.03±1.58 | 11.89±0.34 | 18.04±5.06 |
| 20% | 2.92±0.25 | 5.13±1.00 | 14.94±0.43 | 20.75±2.84 |

Similarly, Table 3 presents the regression analysis of predicted versus FEM reference temperatures across different input noise levels. For instance, under noise-free conditions (noise level = 0), the model achieves a slope of 1.0004 ± 0.0026, an intercept of 0.0000 ± 0.0001, and an R² score of 0.9997 ± 0.0002 for the FullFire beam, and a slope of 0.9969 ± 0.0104, an intercept of −0.0002 ± 0.0005, and an R² score of 0.9931 ± 0.0047 for the SideFire beam, indicating near-perfect linear agreement and unbiased prediction performance in both laser beam types. When the noise level increases further to 20%, the slope, intercept and R² score have little changes for both FullFire and SideFire beams, reflecting the minor effect of noise perturbations on the model’s predictive linearity.

**Table 3.** Regression scores between the ground truth and prediction.

| Noise level | Slope | Intercept | R<sup>2</sup> score |
|---|---|---|---|
| FullFire |  |  |  |
| 0 | 1.0004±0.0026 | -0.0000±0.0001 | 0.9997±0.0002 |
| 5% | 0.9999±0.0016 | -0.0000±0.0001 | 0.9997±0.0001 |
| 10% | 1.0009±0.0027 | 0.0000±0.0001 | 0.9994±0.0002 |
| 15% | 1.0008±0.0029 | -0.0000±0.0001 | 0.9991±0.0002 |
| 20% | 1.0001±0.0039 | 0.0000±0.0001 | 0.9986±0.0003 |
| SideFire |  |  |  |
| 0 | 0.9969±0.0104 | 0.0002±0.0005 | 0.9931±0.0047 |
| 5% | 0.9995±0.0085 | -0.0000±0.0004 | 0.9948±0.0065 |
| 10% | 0.9979±0.0074 | 0.0001±0.0003 | 0.9960±0.0030 |
| 15% | 0.9999±0.0059 | 0.0001±0.0002 | 0.9970±0.0009 |
| 20% | 0.9967±0.0084 | 0.0002±0.0003 | 0.9940±0.0027 |

### 3.3 Prediction Error of Thermal Dose Distribution

The trained DeepoLITT model was also applied to infer the dynamic temperature evolution and corresponding CEM43 thermal dose distributions. Figures 8 and 9 illustrate two representative synthetic LITT procedures performed with FullFire and SideFire laser beams under typical clinical settings (FullFire: power = 10 W, duration = 160 s; SideFire: power = 12 W, duration = 64 s). Temperature fields were computed at each time frame using both the FEM and DeepoLITT methods for comparison. Corresponding CEM43 isodose contours at thresholds of 2, 10, and 60 minutes were subsequently derived.

**Figure 8.**
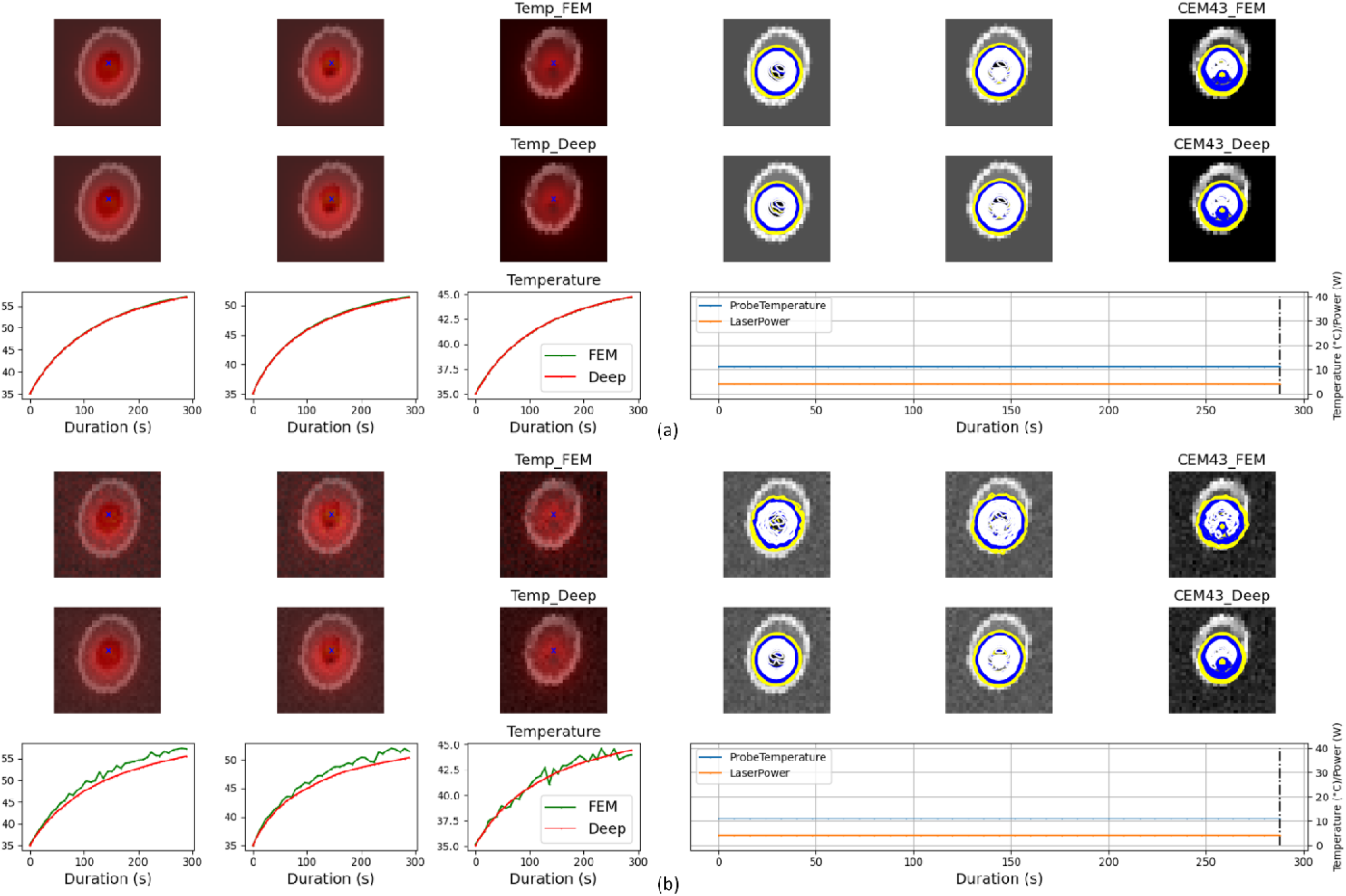
Predicted thermal dose distributions and temporal courses for a FullFire laser beam. (a) at noise level = 0. (b) at noise level = 20%. Temp_FEM and Temp_Deep denote temperature maps generated from the FEM method (ground truth) and predicted from the Deep (the proposed method), respectively. The temporal curves depict time courses of temperature from the two methods at a location indicated by “×” in the images. Contours with yellow, blue and white colors denote convoluted CEM43 iso-doses at 2, 10, and 60 minutes along time course from the two methods.

**Figure 9.**
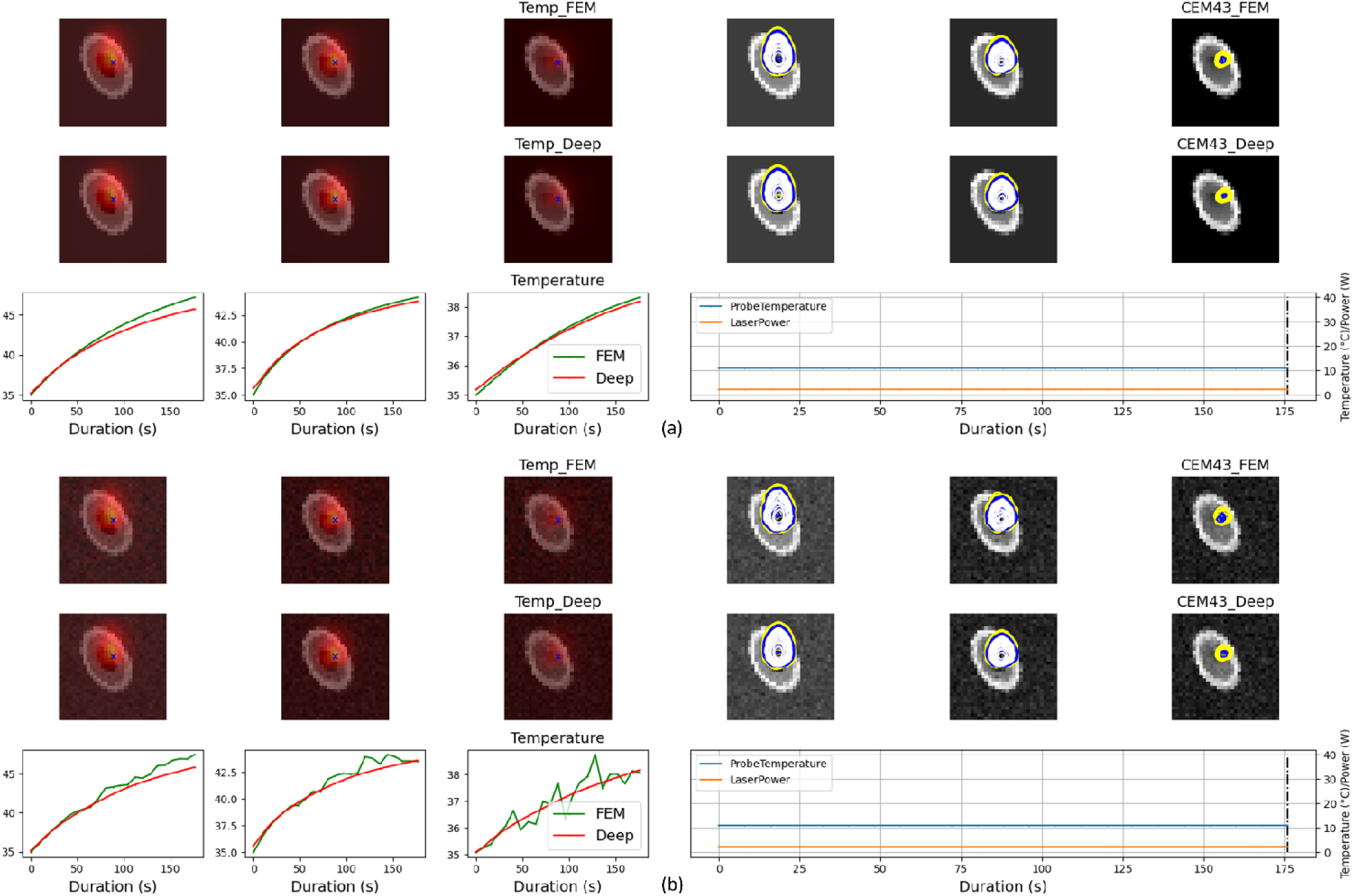
Predicted thermal dose distributions and temporal courses for a SideFire laser beam. (a) at noise level = 0. (b) at noise level = 20%. Temp_FEM and Temp_Deep denote temperature maps generated from the FEM method (ground truth) and predicted from the Deep (the proposed method), respectively. The temporal curves depict time courses of temperature from the two methods at a location indicated by “×” in the images. Contours with yellow, blue and white colors denote convoluted CEM43 iso-doses at 2, 10, and 60 minutes along time course from the two methods.

Visually, the CEM43 contour patterns predicted by DeepoLITT at different noise levels show strong agreement with the FEM reference results across all dose thresholds, as demonstrated in the figures. Quantitatively, for the FullFire beam as tabulated in Table 4, the DSCs averaged across time at zero noise level are 0.96±0.02, 0.97±0.21, and 0.96±0.21, with corresponding HDs of 0.09±0.08 mm, 0.07±0.07 mm, and 0.06±0.08 mm for the 2-, 10-, and 60-minute dose contours, respectively. For the SideFire beam, the DSCs are 0.95±0.18, 0.95±0.17, and 0.91±0.3, and the HDs are 0.20±0.31 mm, 0.11±0.30 mm, and 0.15±0.31 mm for the 2-, 10-, and 60-minute dose contours, respectively. As the noise level increases, the DSC values show a decreasing trend and the HD values increase, as expected. At the highest noise level (20%), the DSCs remain above 0.91 for the FullFire probe and 0.84 for the SideFire probe, while the HDs stay below 0.32 mm and 0.40 mm, respectively. These consistently high DSCs and low HDs across all dose levels and noise conditions demonstrate the model’s strong spatial agreement with FEM references and its robustness in predicting accurate thermal dose distributions under varying procedural conditions.

**Table 4.** DSCs and HDs of CEM43 contours between the prediction and the ground truth at different noise levels.

| Noise Levels | CEM43=2 |  | CEM43=10 |  | CEM43= 60 |  |
|---|---|---|---|---|---|---|
|  | DSC | HD (mm) | DSC | HD (mm) | DSC | HD (mm) |
| FullFire |  |  |  |  |  |  |
| 0 | 0.96±0.02 | 0.09±0.08 | 0.97±0.21 | 0.07±0.07 | 0.96±0.21 | 0.06±0.08 |
| 5% | 0.95±0.03 | 0.10±0.09 | 0.94±0.09 | 0.12±0.18 | 0.94±0.07 | 0.10±0.17 |
| 10% | 0.94±0.02 | 0.15±0.12 | 0.93±0.09 | 0.17±0.18 | 0.96±0.08 | 0.14±0.17 |
| 15% | 0.93±0.05 | 0.24±0.18 | 0.94±0.12 | 0.24±0.24 | 0.95±0.08 | 0.21±0.24 |
| 20% | 0.91±0.21 | 0.32±0.24 | 0.93±0.13 | 0.32±0.30 | 0.94±0.06 | 0.25±0.25 |
| SideFire |  |  |  |  |  |  |
| 0 | 0.95±0.18 | 0.20±0.31 | 0.95±0.17 | 0.11±0.30 | 0.91±0.3 | 0.15±0.31 |
| 5% | 0.93±0.14 | 0.20±0.29 | 0.93±0.08 | 0.13±0.18 | 0.93±0.16 | 0.18±0.33 |
| 10% | 0.93±0.12 | 0.23±0.28 | 0.94±0.10 | 0.14±0.24 | 0.93±0.13 | 0.18±0.29 |
| 15% | 0.90±0.06 | 0.26±0.18 | 0.90±0.10 | 0.18±0.22 | 0.90±0.05 | 0.19±0.19 |
| 20% | 0.88±0.24 | 0.40±0.69 | 0.85±0.17 | 0.23±0.38 | 0.84±0.19 | 0.27±0.54 |

In terms of computational speed, the DeepoLITT model achieved an inference speed of approximately 0.04±0.05 s per frame, compared with 24.32±2.12 s for FEM on an image dimension of [31 × 31 × 3] from 20 cases, representing a speedup of more than two orders of magnitude.

## 4. Discussions

By incorporating LITT procedure parameters (tumor images, laser parameters, and spatial coordinates) into a neural network framework, this study proposes a method to rapidly predict heat spread in LITT procedures. After being trained and tested using synthetic datasets, the model achieved promising results: as the input-noise level increases from 0 to 20%, the average temperature prediction errors range from 1.16% to 2.92% for the FullFire laser probe, and 3.79 % to 5.13 % for the SideFire laser probe. For the thermal dose prediction, the model showed a strong agreement with the FEM-based ground truth, achieving DSCs from 0.97 to 0.91 (FullFire) and from 0.95 to 0.84 (SideFire), and HDs from 0.06 to 0.32 mm (FullFire) and from 0.11 to 0.40 mm (SideFire). In terms of computational efficiency, the proposed DeepoLITT framework achieved an inference time of 0.04 s per 3D sample ([31, 31, 3]), compared to 5.04 s for the FEM method, representing more than a 100× speedup while maintaining comparable accuracy. These results collectively highlight the model’s accuracy, precision, robustness, and computational efficiency, underscoring its suitability for rapid and reliable thermal dose prediction. The promising performance obtained on synthetic datasets lays the groundwork for our future investigations using clinical data.

The proposed neural network builds upon the DeepONet architecture, which was originally introduced by Lu et al.^19,20^. Since proposed, the DeepONets have been successfully applied in resolving complex physics problems including thermal ^22,28^, optical ^45^, energy^46^, and multi-physics ^24,47,48^. The essence of these DeepONet implementations depend on resolving a functional mapping problem between input and output functions ^19,20^. In the context of LITT, the operator can be conceptualized as a mapping from tumor tissue properties or laser source profiles to the resulting spatiotemporal temperature distributions. However, in real clinical practice, rigorous characterization of in vivo tumor tissue properties and in situ laser source profiles is highly challenging. For instance, although optical and thermal properties of ex vivo tumor tissues have been reported^34–39^, corresponding in vivo measurements remain largely undocumented. Likewise, direct assessment of laser source profiles is difficult because the structural and material specifications of commercial laser probes are proprietary and generally inaccessible to investigators. To address these limitations, we propose an alternative strategy that incorporates readily available global laser parameters—such as power, duration, and probe orientation—together with tumor imaging into the functional mapping. These inputs are routinely collected in clinical workflows. For example, laser power, duration, and probe orientation are prescribed by surgeon before each LITT procedure, and pretreatment MRI images of brain tumor patients are routinely scanned and archived in PACS or BrainLab systems. By leveraging this novel formulation and clinically accessible information, the proposed model could be feasibly deployed in real-world clinical environments.

We did not incorporate explicit physics equations into the model to construct a physics-informed neural network, although this approach has shown potential for reducing overfitting and improving generalization^18,49,50^. However, prior work has also reported that physics-informed DeepONets can, in some cases, degrade prediction accuracy and introduce additional computational burden^50^. In our study, the complex tumor microenvironment and the lack of documented in vivo tumor properties make it unlikely that classical PDE formulations can fully or accurately characterize heat transport in tumor tissue. Embedding such uncertain or incomplete physics into the model could compromise its ability to learn the underlying functional mapping and, consequently, impair temperature prediction performance.

Regarding the network architecture, we adopt the trunk–branch framework with two branch networks and constructed the model using a combination of convolutional and residual networks. The original DeepONet formulation does not prescribe the number or type of branch networks, nor does it specify how they should be integrated; instead, the authors provide a fully connected network (FCN) only as an illustrative example ^19,20^. In fact, the DeepONet framework explicitly encourages the use of alternative architectures, including convolutional neural networks (CNNs) and attention-based mechanisms such as transformers. A variety of network architectures have since been explored by other investigators, including CNNs^51^, residual networks^22^, U-Nets^22^, gated recurrent units (GRUs) ^52^, long short-term memory (LSTM) networks^52^. Compared with standard FCNs, residual networks are well known for mitigating the degradation problem in deep architectures^30^, while CNNs remain the established approach for extracting informative features from image-based inputs^31^. In this study, we therefore designed our branch and trunk networks using convolutional and residual blocks to leverage their complementary strengths.

To evaluate the contribution of key components to overall model performance, we conducted a series of ablation studies. Hyperparameter optimization was also performed to identify the optimal architectural dimensions and network depth. Rather than manually testing each model variant, a process that is both labor-intensive and computationally costly, we employed the automated Optuna framework^32,33^, which efficiently explores model configurations and tunes hyperparameters. This approach also quantifies the relative importance of individual components and hyperparameters, providing valuable guidance for efficient fine-tuning in future experiments.

Notably, the optimized configuration yields a streamlined architecture with fewer and smaller branch networks, while maintaining strong predictive performance. Training under this optimized setting requires only 3–4 hours for the full 5-fold cross-validation ensemble, representing a substantial reduction in computational cost compared with training larger, manually designed architectures.

To prevent overfitting and enhance generalization, we incorporated several strategies into the training pipeline. Data augmentation (N = 400 samples) was used to increase dataset diversity, and dropout was applied during training to encourage the network to learn redundant and more robust features. Regularization terms were added to the loss function to penalize large weights (weight decay coefficient), and early stopping was employed to halt training when the validation loss no longer improves. A learning-rate scheduler was used to avoid local minima and promote faster convergence. In addition, layer normalization was implemented to mitigate internal covariate shift, stabilize training, and reduce sensitivity to initialization and noise. Finally, a deterministic L-BFGS optimizer was applied after stochastic optimization to further refine model parameters and reduce prediction uncertainty.

We adopted a cross-validation ensemble method for model training, validation and inferencing. Compared with standard cross-validation or standalone ensemble methods, this hybrid strategy integrates the strengths of both approaches. It benefits from the robust training afforded by multiple data splits in cross-validation and the variance reduction achieved through ensembling ^53,54^.

To evaluate reliability of the proposed DeepoLITT model under measurement uncertainties, a noise perturbation analysis was conducted by introducing Gaussian noise into the input datasets. This approach simulates variations commonly observed in experimental or clinical data, such as noise in MR image acquisition and temperature measurement. The results demonstrate that the DeepoLITT maintains high predictive accuracy, low bias, high linearity, and consistently high R² values even at high noise levels, with only minor deviations in temperature fields and thermal dose contours, as evidenced by high DSCs and low HDs in the evaluation. Possible explanations may be due to model’s inherent operator-learning structure which can capture underlying functional relationships rather than memorizing specific data patterns, making it intrinsically more robust to perturbations compared to conventional DL architectures.

Although our study achieved promising results with high predictive accuracy and low uncertainty, the peak L1 errors from those evaluations remain relatively high, particularly for the SideFire laser probes. A likely contributing factor is the limited angular resolution of the laser probe orientations represented in the dataset, which may not sufficiently capture the full variability of angular effects. Increasing the number of randomized augmentations could help alleviate this issue, but doing so would introduce additional computational burden.

The application of deep learning methods to model temperature distribution and thermal dose in MRI-guided LITT and related neuro-ablative procedures remains relatively limited in the literature. Sadatamin et al. employs U-Net ^25^and LSTM ^27^ architectures to predict spatiotemporal temperature evolution in mesial temporal lobe epilepsy (MTLE) lesions, demonstrating the feasibility of deep learning for LITT temperature field estimation but relying primarily on image-to-image mapping between pre-operative MRI and temperature images. Gao et al. further utilize a ConvLSTM ^26^ network to predict thermal damage, represented by the enhancing rim on post-contrast T1-weighted MRI, from intraoperative MR thermometry time-series data. While these approaches show promising accuracy within constrained experimental conditions, their dependence on direct image mappings inherently limits adaptability to changes in procedural settings, such as variations in laser power, exposure duration, or probe placement. In contrast, our proposed DeepoLITT framework advances beyond conventional voxel-level mappings by explicitly incorporating LITT procedural parameters into the learning process. This design enables the model to predict temperature and thermal dose under diverse treatment conditions, supporting personalized and flexible simulation of LITT procedures. Importantly, we replicate identical experimental setups to our clinical LITT procedures, including spatial dimensions, image slice configurations, laser probe specifications, and laser power settings. This strategy facilitates efficient transfer learning and fine-tuning using limited clinical datasets in the next phase of our investigation, thereby bridging the gap between synthetic modeling and real patient-specific data for accurate and reliable thermal dose prediction.

## Limitations

Despite these promising results, limitations of the current study warrant attention in future work:

1). In this study, synthetic data were used to approximate tumor tissue structure and thermal behavior during the LITT procedure. These synthetic representations necessarily oversimplify complex biological features such as tumor heterogeneity, necrotic patterns, microvasculature, and irregular tumor boundaries. Moreover, the simulated images and temperature fields don’t reproduce real-world MRI artifacts or MRT thermometry characteristics, including bias fields, motion, coil inhomogeneity, and probe-induced susceptibility effects. As a result, models trained solely on synthetic data may learn features that do not fully generalize to clinical datasets. Ongoing work is addressing this limitation by incorporating clinical patient images and MRT temperature maps into model training.

2). Single-channel images were used as inputs to encode tumor features in this study. This limitation restricts the model’s ability to represent the full biological and structural heterogeneity of brain tumors. Future work involving clinical data will incorporate multimodal MRI inputs such as T1-weighted, T2-weighted, and diffusion-weighted images to more comprehensively characterize tumor environments and thereby improve the accuracy of thermal dose prediction.

## 5. Conclusions

In this study, we propose DeepoLITT, a deep neural operator designed to deliver accurate, rapid, and robust predictions of temperature and thermal dose distributions in MRI-guided LITT procedures for brain tumors. By formulating a functional mapping that integrates laser parameters with tumor imaging data and constructing a trunk–branch architecture built on convolutional and residual networks tailored for LITT procedure parameters, we developed a model capable of predicting temperature fields and thermal dose distributions to support treatment planning.

Leveraging the hyperparameter optimization framework, we perform automated architecture ablation and hyperparameter optimization to refine and simplify the model structure. The resulting model demonstrates strong agreement with FEM references, achieving low temperature prediction errors and high thermal dose similarity across a range of noise levels. In terms of computational efficiency, DeepoLITT achieves more than a 100× speedup over conventional FEM solvers while maintaining comparable accuracy, underscoring its potential for rapid preoperative thermal dose prediction.

Future work will incorporate clinical patient data to further validate and refine the model.

